# Perceptual decision-making: extending a standardized rodent task to humans

**DOI:** 10.64898/2026.10.01.755923

**Authors:** Isabelle Hoxha, Philippa A. Johnson, Fenying Zang, Camilla Ucheoma Enwereuzor, Anne E. Urai

## Abstract

Mice are widely used to study the neural basis of perceptual decision-making, often with the ultimate goal of extrapolating insights to humans. Yet, whether humans and mice rely on similar computational strategies remains rarely tested, partly because the two species acquire knowledge differently. Animals learn from experience, while humans usually receive verbal task instructions. Here, we explicitly contrast these modes of learning in humans who perform a perceptual decision-making task adapted from a standardized rodent task. Specifically, we present an open behavioral dataset from 80 human participants who performed 600 trials of a visual contrast-discrimination task, matched to the standardized International Brain Laboratory task for mice (International Brain Laboratory et al., 2021, 2025). Participants either received verbal instructions (‘Instructed’) or were only told to discover the rules of the task by playing (‘Uninstructed’). The dataset contains choices, response times, continuous cursor trajectories, eye-tracking and pupil recordings. Descriptive analyses show that Uninstructed participants were less accurate and had higher lapse rates than Instructed participants. They improved over the first 200 trials, approaching the performance of Instructed participants. These data provide a benchmark for testing whether computational strategies identified in mice also describe human behavior.

## Introduction

Rodents, and especially mice, have become a leading model for studying the neural basis of perceptual decision-making because they can be trained to perform quantifiable psychophysical tasks while neural activity is measured at scale (Carandini & Churchland, 2013; Hanks & Summerfield, 2017; International Brain Laboratory et al., 2025; Urai et al., 2022). A central goal of this work is to uncover principles of decision-making that apply across species, and particularly translate to humans. Whether insights transfer across species, however, relies on a premise that is rarely tested directly: that different species solve similar tasks using the same computational strategies (Barron et al., 2021; Chakravarty et al., 2025; Glukhova et al., 2026; Musall et al., 2019). A quantitative characterization of behavior is a prerequisite for validating this premise (Gomez-Marin & Ghazanfar, 2019; Krakauer et al., 2017).

Matched comparisons of humans and rodents are therefore valuable, and have revealed both similarities and differences. Rats and humans performing the same auditory task accumulate evidence near optimally (Brunton et al., 2013). Mice (Fernandez et al., 2025; Findling et al., 2025), rats (Akrami et al., 2018) and humans (Urai et al., 2019) show sequential effects of past stimuli and choices. Computational models have demonstrated that rodents and humans use divergent strategies in visual object recognition (Schnell et al., 2023). Such direct comparisons remain rare, and openly shared datasets of humans and rodents performing the same task are scarce, although large datasets exist for each species (Bonacchi et al., 2023; Glasser et al., 2016; International Brain Laboratory et al., 2025; Rahnev et al., 2020).

In perceptual decision-making experiments, observers can learn about the structure of the task in different ways: through instruction or experience. This complicates cross-species comparisons. In non-human animals, task knowledge is acquired from experience (usually guided by very specific protocols to lead animals to task knowledge and decision strategies), while humans are usually told what to do. Verbal instructions allow people to perform a new task after a single exposure (Cole et al., 2013) and can bias or even override learning from experience (Li et al., 2011). Even when overall performance is similar, learning from description and learning from experience can lead to systematically different choice patterns (Garcia et al., 2021; Hertwig & Erev, 2009), and to different use of prior information in perceptual inference (Bévalot & Meyniel, 2024; Forsgren et al., 2025).

Here, we collected an open dataset of 80 human participants who performed a visual contrast-discrimination task adapted from the standardized mouse task of the International Brain Laboratory (International Brain Laboratory et al., 2021). We preserved key elements of this original task: a continuous motor response, uncued blocks with changing stimulus probabilities, and identical stimulus sequences. We manipulated how participants acquired task knowledge by creating two experimental groups who either did or did not receive verbal instructions on the stimulus-response mapping of the task. In addition to choices and response times, we recorded continuous cursor trajectories and pupillometric measures, as well as facial video. This paper documents the dataset and reports descriptive analyses of choice behavior.

## Results

Human participants (n=80, 65 female, 13 male, and 2 nonbinary participants; mean age = 19.99 years, SD = 3.14, range = 18–35 years) completed a perceptual decision making task, designed to be similar to a standardized perceptual decision-making task designed for mice, hereafter called ‘the IBL task’ (International Brain Laboratory et al., 2021) (Figure 1Aa). In each trial, participants observed two gratings presented on the screen and decided which had the higher contrast (target stimulus). They indicated this by moving the computer mouse (hereafter called the ‘cursor’) to bring the target stimulus into the centre of the screen (Figure 1Ab). Each participant was assigned to one of two conditions: one in which participants were instructed about the stimulus-reward contingency and how to make responses (Instructed group, n=39, 32 female, 6 male, and 1 nonbinary participants; mean age = 19.74 years, SD = 3.08, range = 18–35 years) and one in which they were only told to respond by moving the cursor but must figure out the rules of the game by playing (Uninstructed group, n=41, 33 female, 7 male, and 1 nonbinary participants; mean age = 20.22 years, SD = 3.23, range = 18–33 years) (Figure 1Ab).

### Uninstructed participants are more prone to lapses than instructed participants

Participant performance was assessed on the basis of the accuracy of their choices in easy trials (i.e. contrast differences >= 50%) (Figure 1C). Uninstructed participants were less accurate than Instructed participants (mean Uninstructed: 83.43% [51.68; 97.65], mean Instructed: 97.26% [92.91; 100], Welch independent t-test: *t*(40.98) = 5.520, *p* < 0.001, *d* = 1.22). Participant behavior at each contrast level was assessed using a four-parameter psychometric curve (see Methods; Figure 1B). Participants in the Uninstructed group had higher lapse rates (mean lapse (Uninstructed - Instructed): 0.12 − 0.01, *t*(40) = 4.46, *p* < 0.001, *d* = 0.98) than Instructed participants. We observed no difference in other psychometric parameters between the two groups (see Table 1).

**Figure 1.**
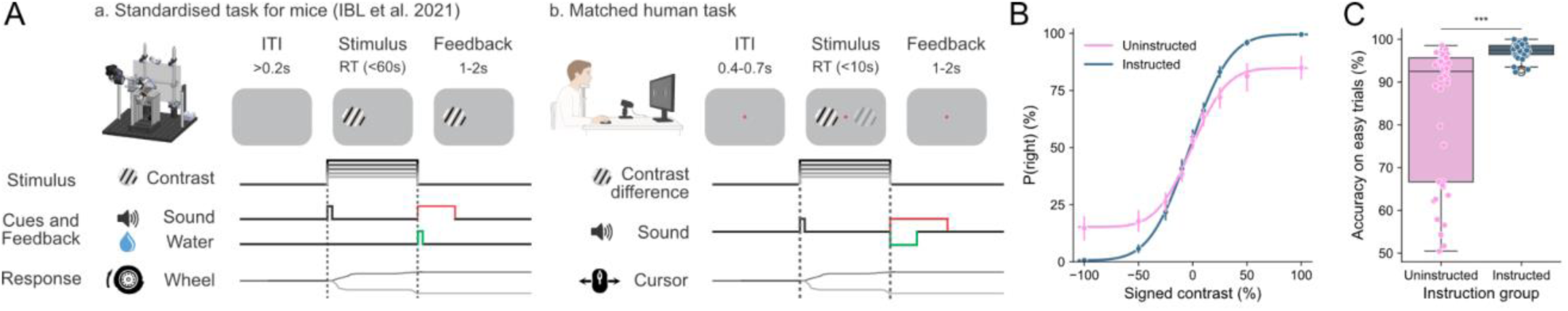
Human task and behavior. **(A)** Task schematic, comparing the IBL mouse task with our human version. Panels adapted from (International Brain Laboratory et al., 2021) under a CC-BY license and from (Soric, 2026) with permission. See also Table 2 for a detailed comparison of the two tasks. **(B)** Psychometric curves for the Instructed and Uninstructed group. Points show the average proportion of rightward choices at each signed contrast across participants (error bars are the 95% confidence intervals (CIs)), while solid lines show psychometric fits to the pooled trial data of each group. **(C)** Performance on easy trials (50% and 100% contrast difference) for the Instructed and Uninstructed group. *** p < 0.001.

**Table 1.** Psychometric parameter summary and statistical tests between the two groups. Absolute bias, slope and mean lapse are computed from psychometric curve fits over each participant’s full trial sequence. Significant differences are in bold.

| Parameter | Uninstructed | Instructed | Statistical test on group difference |
| --- | --- | --- | --- |
| Absolute bias | 7.16<br>[5.14, 9.18] | 6.00<br>[4.22, 7.78] | $t(77.2) = 0.87, p = 0.387, d = 0.19$ |
| Slope | 32.65<br>[26.03, 39.26] | 38.71<br>[36.29, 41.13] | $t(50.4) = 1.74, p = 0.088, d = 0.39$ |
| Mean lapse | 0.12<br>[0.07, 0.18] | 0.01<br>[0.00, 0.01] | $t(40.2) = \mathbf{4.46}, p < \mathbf{.001}, d = \mathbf{0.98}$ |

**Table 2.** Key differences between the IBL task and the human task (see also Figure 1A).

| Task feature | IBL mouse task (IBL et al. 2021) | Human task |
| --- | --- | --- |
| Task | Detection of grating from 0-100% contrast | Discrimination of target grating (50-70% contrast) from non-target grating (50% contrast) |
| Response modality | Wheel | Computer mouse (cursor) |
| Response time out | 60 s | 10 s |
| Feedback | Correct: sugar water. Error: white noise + 1 s time-out | Correct: high beep. Error: white noise + 1 s time-out |
| Session end condition | Any of three criteria:<br>(1) <400 trials in 45 min;<br>(2) >400 trials and session duration ≥ 90 min; or<br>(3) >400 trials and slow-down (median RT over the last 20 trials $>5\times$ the whole-session median) | 600 trials completed |

### Block-dependency and statistical learning

As in the original IBL task, our task included uncued blocks where the probability of the target stimulus appearing on the left was 50% (first 90 trials of the session), 80% or 20% (variable block length over the rest of the session) (Figure 2A). In mice, blocks were introduced only after animals gained a basic performance on the task (International Brain Laboratory et al., 2021), and the effect of this block structure on choice behavior grew over sessions (Roy et al., 2021).

**Figure 2.**
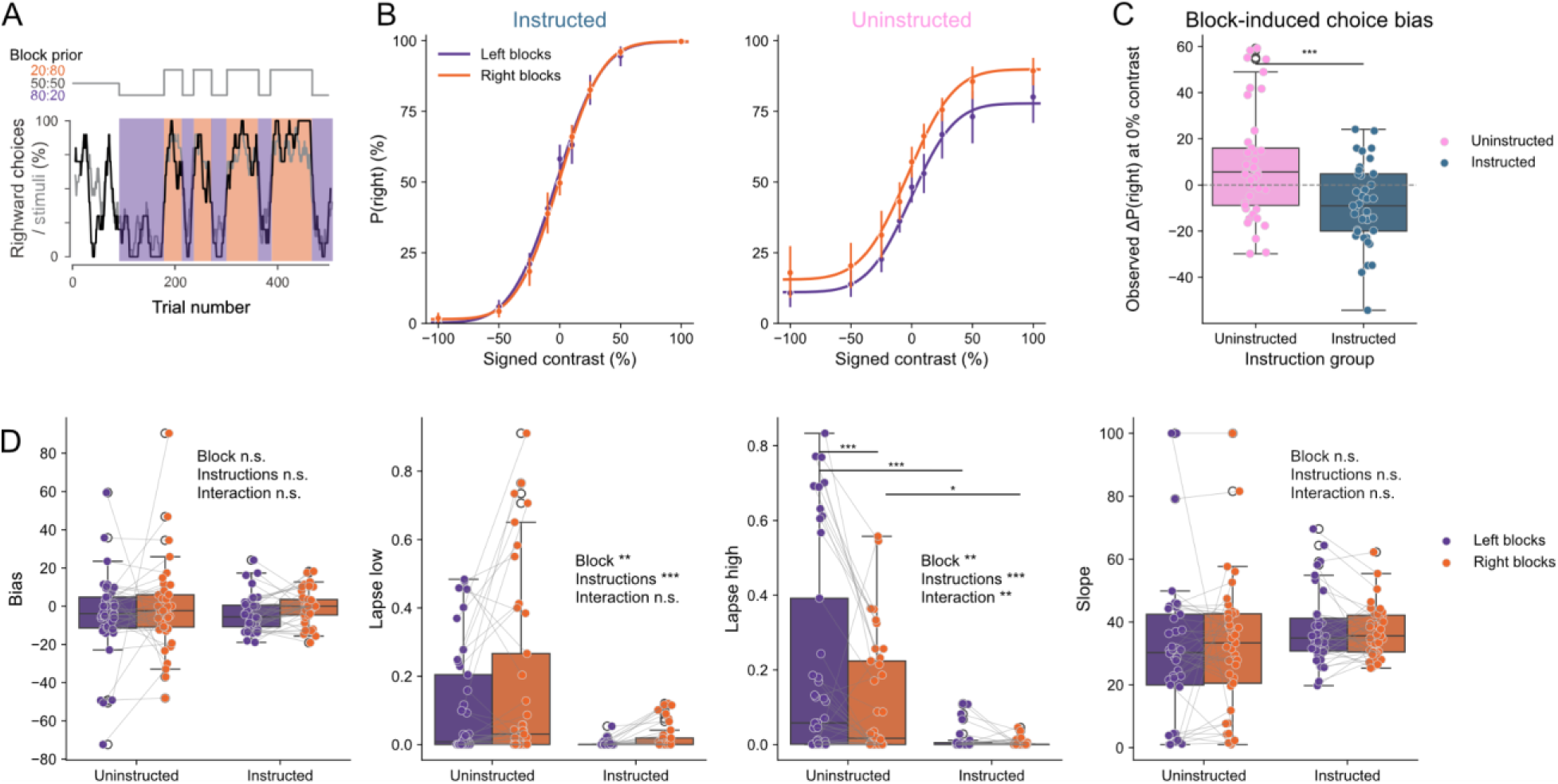
Behavioral performance and block-induced bias. **(A)** Block structure in the task. After the first 90 trials, in which the target stimulus had 50% probability of appearing on the left, the probability of the target stimulus appearing on the left switched uncued between 20% and 80%. **(B)** Psychometric curves conditioned on block type for Instructed (left) and Uninstructed (right) participants. Instructed participants received explicit instructions about stimulus-reward contingencies but not about the block structure. Points show the average proportion of rightward choices at each signed contrast across participants (error bars are the 95% CIs), while solid lines show psychometric fits to the pooled data. Purple and orange indicate left- and right-biased blocks, respectively. **(C)** Block-induced choice bias on 0% signed contrast (contrast differences = 0%) trials, calculated for each participant as ΔP(right) = P(right | right block) − P(right | left block). Points represent individual participants, bars show group means, and error bars indicate 95% CIs. Positive values indicate a greater probability of choosing right in right-biased than in left-biased blocks. **(D)** Comparison of psychometric curve parameters (signed bias, lapses and slope) between blocks for Instructed and Uninstructed participants. Each dot represents a fitted value for one participant, and the grey lines link parameter pairs of one participant. Only the interaction post-hoc tests are displayed as bars on the panel. ∗ p < 0.05,∗∗ p < 0.01,∗∗∗ p < 0.001.

In our human dataset, the effects of this block manipulation varied between the Instructed and Uninstructed groups (Figure 2B). The block effect on choice at zero contrast was significant in both groups (paired-samples t-test, Uninstructed: *t*(40) = 2.369, *p* = 0.02, Instructed: *t*(38) = 3.035, *p* = 0.004), and this block-dependent choice at zero contrast shift was larger in the Uninstructed group than in the Instructed group (Welch t-test, *t*(73.04) = 3.714, *p* < 0.001, *d* = 0.770) (Figure 2C).

We next quantified these block effects by fitting psychometric functions conditioned on the block, and comparing the four parameter values (Figure 2D). For the low lapse (i.e. probability of making a random choice on a high left contrast stimulus), we observed a significant main effect of block type (*F*(1,78) = 7.63, *p* = 0.007), a main effect of instructions (*F*(1,78) = 15.90, *p* < 0.001) on the low lapse, Post-hoc analyses show that there were fewer low lapses on left-biased blocks than right-biased blocks (mean difference±standard error: −0.042 ± 0.015, *t* = 2.762, *p_Holm_* = 0.007) and higher for uninstructed participants than instructed participants (mean difference: 0.129 ± 0.032, *t* = 3.987, *p_Holm_* < 0.001). Similarly, for the high lapse (i.e. probability of making a random choice on a high right contrast stimulus) we observed a main effect of block type (*F* = 10.57, *p* = 0.002), a main effect of instructions (*F*(1,78) = 22.35, *p* < 0.001), and an interaction effect (*F*(1,78) = 7.89, *p* = 0.006). Post-hoc analyses reveal that high lapse rates were higher in left-biased than right-biased blocks for Uninstructed participants (0.101 ± 0.023, *t* = 4.339, *p_Holm_* < 0.001), whereas no significant between-block difference was detected for Instructed participants (0.007 ± 0.024, *t* = 0.308, *p_Holm_* = 0.759). High lapse rates were also higher for Uninstructed than Instructed participants in both left-biased blocks (0.205 ± 0.037, *t* = 5.484, *p_Holm_* < 0.001) and right-biased blocks ( 0.111 ± 0.037, *t* = 2.975, *p_Holm_* = 0.011; see Supplementary Table 1). Notably, we did not observe significant differences in the slopes (main effect of block: *F*(1,78) = 0.016, *p* = 0.900:, main effect of instruction: *F*(1,78) = 2.728, *p* = 0.103, interaction: *F*(1,78) = 0.252, *p* = 0.617) or bias term (main effect of block: *F*(1,78) = 1.793, *p* = 0.184:, main effect of instruction: *F*(1,78) = 0.052, *p* = 0.821, interaction: *F*(1,78) = 0.008, *p* = 0.927) of the average psychometric function. This latter finding, especially in contrast with the observed group difference in choice bias (Figure 2C), indicates that our two ways of operationalizing choice bias (either directly on choice fractions on only 0% contrast trials, or by estimating the offset from the full psychometric function fits) do not always lead to the same conclusions (see Methods).

In the next sections, we present qualitative descriptions of response times, cursor trajectories and learning dynamics. They are meant to characterize the data rather than test hypotheses, as robust statistical inference will require more in-depth modelling, which we leave for future work.

### Group differences in response times

Response times (RTs) offer another window into participants’ behavioral strategies. An example Uninstructed participant had higher RTs for the first ∼150 trials, and then showed a fairly consistent RT, indicating a period of learning followed by stable task performance. In contrast, the example Instructed participant showed stable RTs throughout the task, with short epochs of longer RTs (Figure 3A). At the group level, the Uninstructed participants had qualitatively similar median RTs (Figure 3D), and more variable RTs over all contrast levels (Figure 3E). Median RTs showed a typical pattern of increasing RT with lower contrast in both groups. There seems to be a similar relationship between contrast and RT variability in the Instructed group, where lower contrast stimuli are associated with increased RT variability, but this pattern is not present in the Uninstructed group.

**Figure 3.**
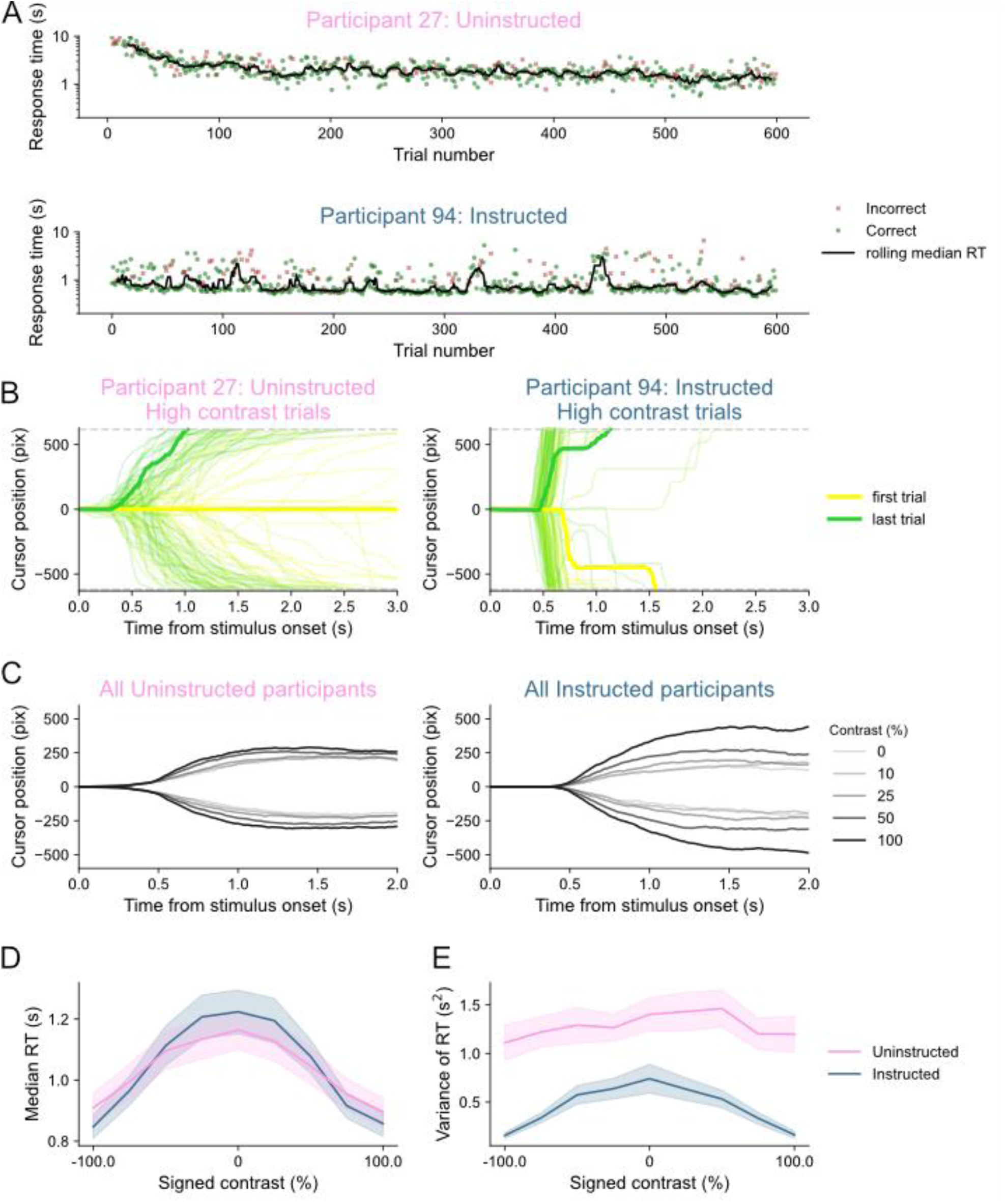
Response times and cursor movements. **(A)** Response times across a single session of responses for an example Uninstructed (top) and Instructed (bottom) participant. Correct and incorrect responses are marked by green points and red crosses, respectively. The black line shows the rolling median RT over 10 trials. **(B)** Cursor trajectories for all high contrast trials of the example Uninstructed (left) and Instructed (right) participant. The first (yellow) and last (green) high contrast trials (100% contrast) of the session are highlighted with thicker lines. Grey dashed lines show the threshold for a left (top) or right (bottom) response. **(C)** Cursor trajectories averaged over all trials with a given contrast level and choice direction for Uninstructed (left) and Instructed (right) participants. **(D)** Median RTs as a function of signed contrast for Instructed and Uninstructed participants. Error bars show 95% CIs. **(E)** Variance of RTs as a function of signed contrast for Instructed and Uninstructed participants. Error bars show 95% CIs.

Cursor trajectories, analogous to the wheel responses in the IBL task (Figure 1A) (Amjad et al., 2026), provide insight into decision formation and execution beyond what is possible in button-press tasks. They allow investigation of movement initiation times, movement speed, and changes of mind (Figure 3B). The example Instructed participant responded with ballistic, faster movements throughout the task, compared to the slower movements of the example Uninstructed participant. Additionally, the Instructed participant made a few slower movements throughout the task, while the Uninstructed participants made slower movements in the early part of the session. Figure 3C shows average cursor trajectories for each group, split by contrast and choice direction. In both groups, higher contrast stimuli seem to result in cursor movements that are initiated earlier. Qualitatively, the Uninstructed group showed less amplitude in average trajectories overall. This could be because they had more incomplete responses that never reached the response threshold. Alternatively, they may have kept the cursor closer to center for longer or used lower movement vigor. Finally, they could show more trial-to-trial temporal variability in when they initiated movements, which would smear the average line. Future investigations could use cursor trajectories to characterize individual differences and within-session variability in decision strategy.

### Group differences are attenuated throughout the experiment but do not disappear

Participant behavior can also be observed over the duration of the experiment (600 trials per participant, without breaks), which allows examination of task dynamics (Figure 3A, 4). We here describe these dynamics qualitatively, acknowledging that full statistical modelling will require more in-depth consideration of the exact learning strategies across individuals.

Participants who received explicit task instructions responded quickly and accurately from the first trials of the session, and their performance changed little over the course of the session (Figures 4A and 4B). Uninstructed participants began at near chance accuracy with longer response times, but improved in both accuracy and response times over the first ∼200 trials. However, at the group level, they did not reach the accuracy of the Instructed participants. We next looked at behavior in three trial windows within each session (Figure 4C): the first 200 trials, trials 200 to 400, and trials 400 to 600. In the first 200 trials, Uninstructed participants’ choices were less sensitive to contrast difference between the two gratings than Instructed participants’ choices, notably on high contrast stimuli. Beyond higher RTs, RT variability was also higher at all contrast levels in early trials. Over the subsequent two windows, the Uninstructed group’s psychometric curves progressively steepened, and their RTs and RT variability decreased, such that by the last trial window the two groups’ curves were largely superimposed. RTs followed an inverted-U shape across contrast in both groups and all trial windows, consistent with trials with small contrast differences being the most difficult trials.

**Figure 4.**
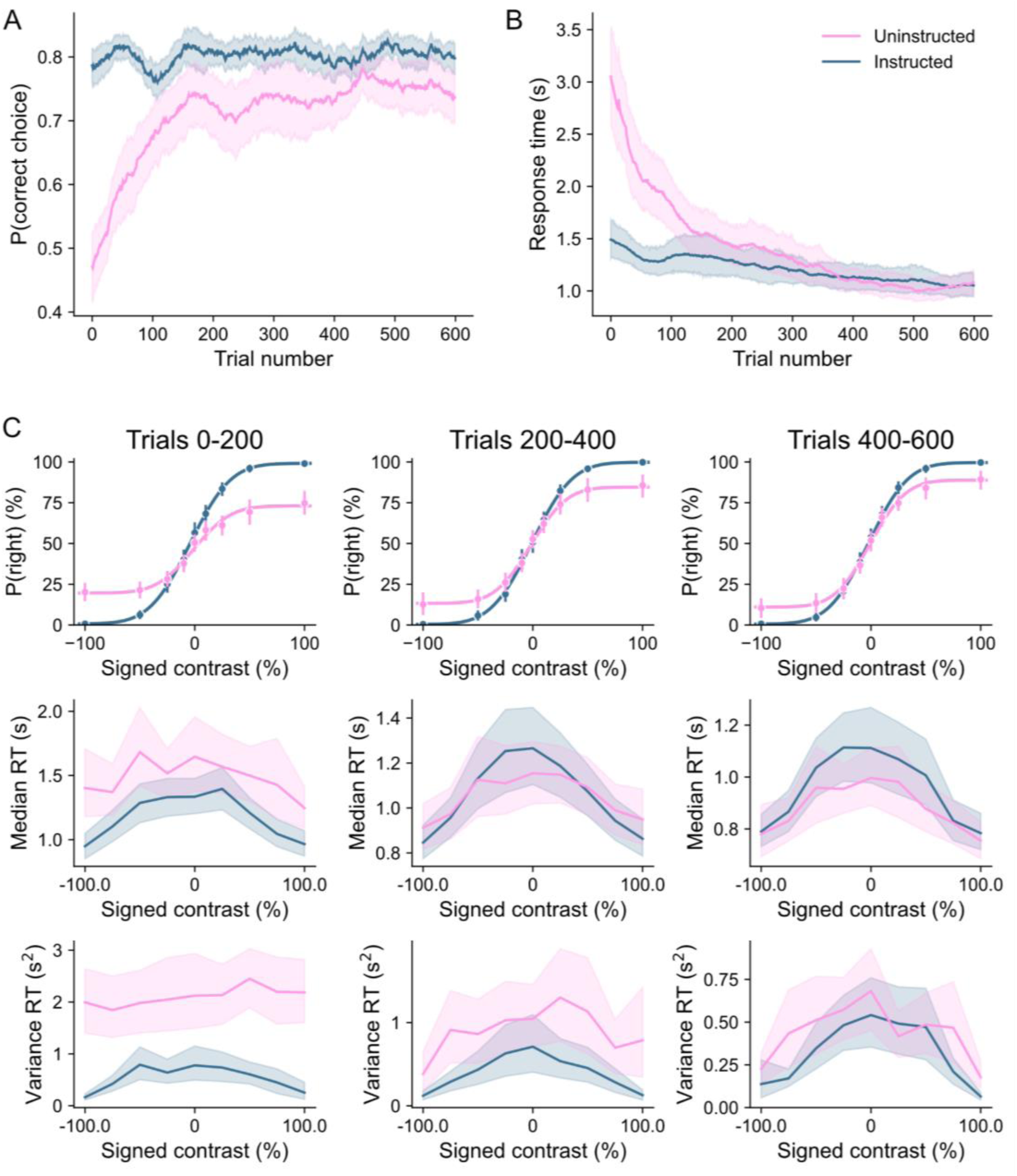
Evolution of behavior over the course of the session. **(A)** Proportion of correct choices as a function of trial number. While the Instructed group’s performance (in blue) stayed stable, the Uninstructed group (in pink) showed an increase in performance over the first ∼200 trials. **(B)** Evolution of median RT as a function of trial number. **(C)** Evolution of psychometric curves (top), median RT (middle) and variance of RT (bottom) across the experiment, computed over three time windows: trials 0-200, 200-400 and 400-600. The two groups differ the most in early trials. Error bars and shaded areas represent 95% CIs over participants.

## Discussion

Here, we build on the large-scale, open methods and data from (International Brain Laboratory et al., 2021, 2025) that provide a benchmark for behavior and neural coding in mice (Andre et al., 2026). To better understand the neural basis of decision-making, it is crucial to explicitly compare the strategies that drive behavior in different species (Urai, 2025). Such comparison presents inherent challenges: an identical task will likely lead to very different behavior, whereas matching for the same behavior may require substantial task adjustments. As in other recent work (Chakravarty et al., 2025; Glukhova et al., 2026), we here find a compromise between those two approaches. We built on a standardized, well-documented task and modified only the elements crucial to obtain similar behavior in humans.

We explored one important difference in comparing humans with other animals: how knowledge of task structure and goals is acquired (Bévalot & Meyniel, 2024). In the large majority of studies on perceptual decision-making in humans, participants receive verbal instructions about the task structure, which allow new tasks to be performed after a single exposure (Cole et al., 2013). Non-human animals instead rely on elaborate training and shaping protocols to build task understanding, which are often very task-specific and of which standardization has only recently been attempted (Burgess et al., 2017; Guo et al., 2014; International Brain Laboratory et al., 2021). In our experiment, we explicitly varied these ways of acquiring task knowledge. Unsurprisingly, all participants who received instructions on the task (but without explicitly being told to watch for block structure or patterns in the target stimulus location) performed well, with a mean accuracy of 97% on easy trials. In contrast, there was considerable variation amongst those who received no instructions, and instead learned the rules of the game by playing (83% average accuracy).

The Uninstructed group specifically provides a window on task learning in a setting that parallels animal training. Learning from description and from experience can lead to systematically different choices, even when overall performance is similar (Hertwig & Erev, 2009), so withholding instructions may bring human behavior closer to that of animals. Matching the method of learning does not, however, guarantee matching learning dynamics. In a recent study of reinforcement learning, humans and rats both valued rewards relative to their context. While humans acquired preferences within a single session, rats needed several, and humans partially corrected their context-dependent biases on transfer trials whereas rats retained them more persistently (Ferguson et al., 2026). Similarly, our Uninstructed participants learned the task within a single session, whereas animals typically require extensive training. Learning from experience thus puts humans and animals on more comparable footing, but differences in learning speed and flexibility can still be expected.

Some participants who did not receive explicit instructions obtained high accuracy (i.e. low number of lapses), showing that they were able to learn the basic stimulus-response mapping of the task. Lapses are often treated as a nuisance, but they can bias estimates of the other psychometric parameters if they are not modelled (Prins, 2012; Wichmann & Hill, 2001). Those with lower accuracy on easy trials may have more complex behavioral strategies that could be dominated by a mix of random responses, choice repetition (Fernandez et al., 2025), and likely include a mix of discrete (Johnson et al., 2025; Nuiten et al., 2026) and slowly drifting (Vloeberghs et al., 2026) behavioral states. In rodents, lapses have been shown to reflect, at least in part, exploration beyond inattention (Ashwood et al., 2022; Pisupati et al., 2021). The post-session debriefing (see Methods) offers a way to relate participants’ reported strategies to their behavior.

The block structure of the task makes it possible to investigate how observers acquire and use prior information. Neither group was informed about the underlying block structure, so any block effect was learned from experience. Humans can adapt to trial-to-trial changes in prior probability (Norton et al., 2019) and in rodents the influence of expectations depends on recent choices (Findling et al., 2025). Our descriptive analyses point to block-related effects that differ between instruction groups, but, as characterizing them properly will require trial-by-trial modelling, we refrain from strong inferences here.

Our task, descriptives and open dataset offer a foundation for more in-depth exploration of specific behavioral strategies - especially amongst those who received no instructions. This could be explicitly linked to behavioral models that have been successfully used on the IBL mouse data: models of prior-based decision-making (Findling et al., 2025), task learning (Bruijns et al., 2026), engagement states (Ashwood et al., 2022; Mohammadi et al., 2025) or slowly fluctuating decision strategies (Roy et al., 2021). Continuous cursor trajectories provide a richer readout of decision formation than button presses, allowing examination of movement initiation (Yin & Hiratani, 2026), vigor (Thura et al., 2025; Thura & Cisek, 2014) and changes of mind (Freeman & Ambady, 2010; Resulaj et al., 2009; Song & Nakayama, 2009; Wispinski et al., 2020). However, direct comparisons with the wheel movements of mice should be made with care, as interactions with a cursor or wheel differ biomechanically in the movement and force required. This dataset also includes pupillometry, eye-tracking and facial video data which can be used to supplement our understanding of the specific computations and strategies used during decision making.

Several limitations should be considered. First, the sample consisted of university students recruited through a participant pool, predominantly female and young, and is not representative of the wider population (Henrich et al., 2010). Second, the Uninstructed condition is only an approximation of animal training, as humans bring extensive prior experience with abstract decision tasks and computer games, receive auditory feedback and a score rather than a primary reward, and learn within a single session instead of over several weeks. Moreover, Instructed participants receive partial instructions and are not informed about the statistical structure of the task. Since the task code is made available, more versions of this task can be collected by other laboratories on other population types to alleviate these limitations.

In conclusion, we present a multimodal extension to humans of a standardized, well-characterized rodent perceptual decision-making task, with a controlled manipulation of the mode of task learning. The dataset and accompanying code are open, documented and aligned to a mouse dataset with identical stimulus sequences. This enables several kinds of reuse: (i) fitting and comparing cognitive models of choice and response times, including latent-state and learning models originally developed on the mouse data (Ashwood et al., 2022; Bruijns et al., 2026; Findling et al., 2025; Mohammadi et al., 2025; Roy et al., 2021), (ii) studying individual differences in how task rules are discovered without instructions (Bévalot & Meyniel, 2024), and (iii) benchmarking cross-species analysis pipelines for cursor kinematics (Amjad et al., 2026) and pupillometry (Johnson et al., 2025) on data with known task structure. These human data enable testing whether computational strategies identified in animals generalize to human behavior, with the in-depth characterization of trial-to-trial variability in decision-making behavior setting the stage for future cross-species comparison of neural data.

## Methods

### Sample

The study was granted ethical approval by the Leiden University Psychology Research Ethics Committee, Leiden, The Netherlands (Ethics no. 2024-09-20-A.E. Urai-V2-5653). Participants were recruited through SONA at the University of Leiden, the Netherlands. 80 participants (65 female, 13 male, and 2 nonbinary participants; mean age = 19.99 years, SD = 3.14, range = 18–35 years) completed the experiment. They were compensated at the standard rates, either monetary or in participant credits.

### Procedure

The experiment began with a brief eye-tracking calibration and validation procedure, then recording of age, gender, and handedness, and several instruction screens (see Appendix). Specifically, participants in the Instructed group received instructions about how to perform the task and the goal of choosing the higher contrast grating (written instructions, example images, and 5 practice trials). The Uninstructed group were told only that their goal was to figure out the rules of the game by themselves and that responses were given by moving the cursor. Each participant then completed one session of 600 trials, without breaks.

### Decision-making task

The human task was designed to retain the key characteristics of the IBL task (International Brain Laboratory et al., 2021), but was adapted to be suitable for human data collection. It was programmed in *PsychoPy* version 2024.1.4 (https://github.com/cocosys-lab/human_IBL_cursor). The goal of the task was to indicate which of two present gratings had higher contrast (the target grating) by moving the cursor to bring the target into the middle of the screen.

Humans would perform at ceiling in detecting all the contrast levels that mice saw, given that the stimulus stays on the screen for the duration of the response period. We therefore changed the human task to a discrimination task, and we changed the set of contrast differences (indicated as ‘signed contrast’ in figures) to span the human psychometric function, based on earlier pilots. Target contrast levels ranged from 0.50 to 0.70 (0.50, 0.52, 0.55, 0.60, 0.70), while the non-target stimulus always had a contrast of 0.50. This makes the human task similar to previous rodent tasks that include a pedestal (Burgess et al., 2017; Steinmetz et al., 2019). The contrast difference was rescaled so that the different levels are displayed as 0, 10, 25, 50 and 100 on the x-axis of the psychometric function.

A red fixation cross was presented in the middle of the screen throughout all trials. Participants were asked to fixate at the central fixation cross throughout the experiment. Each trial consisted of two Gabor patches (gratings) with varying contrasts appearing on the screen, one on the left and one on the right of the fixation point. The gratings had a 618x618 pixel diameter, with a spatial frequency of 0.0073 cycles per pixel and a randomly generated phase. Target gratings could appear on the left of the screen with 50% probability (first 90 trials only), or with 20% or 80% probability, with uncued switches. This was determined from ten [pre-generated sequences](https://github.com/int-brain-lab/iblrig/tree/old_master/pybpod_fixtures/IBL/tasks/_iblrig_tasks_ephysChoiceWorld/sessions) taken from the IBL task. These sequences were shown to mice that each performed multiple recording days. The rationale for these sequences was to allow comparison of the behaviour of different mice to the exact same stimulus sequences. We applied the same rationale in the present experiment: each sequence was seen by approximately four Instructed and four Uninstructed participants.

To make the human task easy to run on standard hardware, responses were given by moving the computer mouse (hereafter called ‘cursor’) instead of a wheel. Both gratings moved with the cursor in a closed loop. The cursor was not visible on the screen. If the middle of the target grating reached the midline of the screen (618 pixels from the central start position), a correct response was recorded; if the middle of the non-target grating reached the centre, an incorrect response was recorded.

Auditory feedback was given upon response completion: for correct responses, there was a high-pitched beep (2,000 Hz; 0.2 s) and, for incorrect responses, a white noise burst (0.5 s). Stimulus onset was accompanied by a sharp beep (5,000 Hz; 0.1 s), and the gratings remained visible until a response was recorded or for a maximum of 10 s. If no response was made within 10 s, the trial was marked as timed out and a low beep sounded (567 Hz; 0.5 s). In the case of incorrect responses or timeouts, a 1s time penalty was added before the next trial.

### Debriefing

Once all trials were completed, participants were asked to answer questions about their strategy and task performance:

1. Please describe how you think the game worked and what you think the rules were.
1.How well do you think you did in the game?
2. What strategies did you use during the game?
3. Did you notice any patterns in the game?

Finally, participants were presented their total score (correct trials/600).

### Psychometric curve

Psychometric curves illustrate how the probability of a rightward choice changes with stimulus position and stimulus contrast. For each session, responses were fitted with a parametric error function using a maximum likelihood procedure:

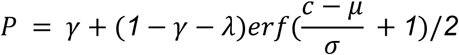

*P* is the probability of rightward choice;

*c* is the contrast level;

*γ*, *λ*, *σ*, *μ* are fitted parameters:

*γ*: low lapse (the lapse rate for left stimuli);

*λ*: high lapse (the lapse rate for right stimuli);

*σ*: slope;

*μ*: the response bias (horizontal shift of the curve)

For details on the fitting process and parameter computation, see (International Brain Laboratory et al., 2021). Note that psychometric fits were not always stable: parameter estimates, particularly for participants with low accuracy where the psychometric function does not describe the data well, could vary considerably between fitting runs, notably because the psychometric parameters are partly redundant (different combinations of bias, slope, and lapse rate can describe the same choice data equally well).

### Statistical analyses

Group differences (Figure 1C, 2C) were evaluated using Welch t-tests, as Instructed and Uninstructed groups had different sizes and had different variances across all metrics. For Welch t-tests, the degrees of freedom are computed using the Welch-Satterthwaite equation, in which the degree of freedom depends on the variance of samples. Therefore, the degrees of freedom are different across statistical tests, unlike in Student’s t-tests, where it only depends on the sample size.

To test for the interaction of group and block effects (Figure 2D), we performed a repeated-measures ANOVA with block as a within-subject factor and instructions group as a between-subject factor. Post-hoc comparisons were done using Holm-corrected t-tests.

### Response time and cursor movement analyses

Response times were measured as the time between stimulus onset and the completion of the response, when the stimulus crossed the midline of the screen. Note that this is after the time of first movement (movement initiation time), which in the rodent task is sometimes called reaction time (Ghani, 2025; Yin & Hiratani, 2026). We quantified response-time variability as the variance of response times across trials within each condition for each participant.

### Eye tracking and pupillometry

Eye-tracking data (gaze and pupillometry data) were recorded using an EyeLink 1000 with chinrest, that was calibrated at the start of each session. See (Johnson et al., 2025) for more details and the pupil data.

### Facial video

Webcam-based video recordings of participants’ faces were obtained during each session, including a mirror placed to capture the computer screen (out of view of the participant). The webcam data was aligned by using computer vision algorithms to detect stimulus onsets in the recorded mirror from the video file (human_ibl_snapshots/utils.py at main · anne-urai/human_ibl_snapshots). These data are not reported here. For privacy reasons, these data can be obtained from the corresponding author on request.

## Code availability

PsychoPy code to run the task is at https://github.com/cocosys-lab/human_IBL_cursor, and all code to analyze the data is at https://github.com/cocosys-lab/2026-human-ibl-analyses. Snapshots of each individual’s performance are at https://github.com/anne-urai/human_ibl_snapshots.

## Data availability

The data are available at https://doi.org/10.5281/zenodo.23072547.

## Contribution statement

Conceptualization: AEU; Data curation: FZ, PAJ; Formal analysis: PAJ, IH, FZ; Funding acquisition: AEU, PAJ, IH; Investigation: CUE (lead); Methodology: CUE (lead), PAJ (support); Software: IH (lead), PAJ, FZ; Supervision: AEU, PAJ (equal); Visualization: FZ, IH, PAJ; Writing – original draft: AEU, IH, PAJ, FZ. Writing – review and editing: all.

## Acknowledgements

This work was supported by the Netherlands Organisation for Scientific Research (Veni fellowship VI.Veni.212.184 to AEU, SSH-XS 406.XS.24.02.032 grant to AEU and PAJ), the Fyssen Foundation (postdoctoral fellowship to IH) and the Chinese Scholarship Council (PhD fellowship 202204910080 to FZ). We thank Annika Frach for coding an earlier version of the task, Maureen Meekel for technical support, and Dorota Moravčíková, Lorenzo van Hoorde, Lucija Sorić, and Cemre Uçkan for assistance with data collection.

## Appendix

Below are the exact instructions given to participants in the Instructed and Uninstructed groups.

### Instructed

You will take part in a computer game.

You will now see some written instructions, as well as two example images of what the stimuli look like. You will then practice the game in a few example trials. Click ‘Continue’ to see the instructions.

On each trial, you must decide which of the two target images has the higher contrast (i.e., appears darker). To select the target with the higher contrast, simply move the mouse until the chosen target reaches the center of the screen. For example, in the image below, the right target has the higher contrast, so you would move your mouse to the left. Click ‘Continue’ to see more instructions and another example.

In this example image, the left target has the higher contrast, so you would move your mouse to the right. To interact with the game, you will only need to move the mouse; clicking does not do anything. Click ‘Continue’ to see more instructions.

You will hear a high beep if you answer correctly. If you answer incorrectly, you will hear a buzzing noise instead. If you take too long to answer, you will hear a low beep. Please try to be as accurate and fast as possible, and try not to let the trial time-out without a response. Click ‘Continue’ to practice a few example trials.

That is the end of the practice trials. Here is a last reminder of the instructions: In each trial, you will hear a beep and two targets will appear. You must decide which target has the higher contrast. To respond, move your mouse left or right to bring the chosen target into the middle of the screen. You don’t need to click, simply drag the mouse. If you are correct, you will hear a high beep; if you are incorrect, you will hear a buzzing noise.

Important: during the game you will see a red cross in the middle of the screen. Please try to keep your eyes fixated on this cross throughout the experiment.

If you have any questions, feel free to ask out loud, the experimenter can hear you and will help you out. You will now start the real experiment. Good luck!

### Uninstructed

You will take part in a computer game.

However, you will not receive any instructions: the goal is to figure out the rules of the game by yourself. To interact with the game, you will only need to move the mouse; clicking does not do anything.

Important: during the game, you will see a red cross in the middle of the screen. Please try to keep your eyes fixated on this cross throughout the experiment.

If you have any questions, feel free to ask out loud, the experimenter can hear you and will help you out. When you are ready, click ‘Start experiment’.

**Supplementary Table 1.**
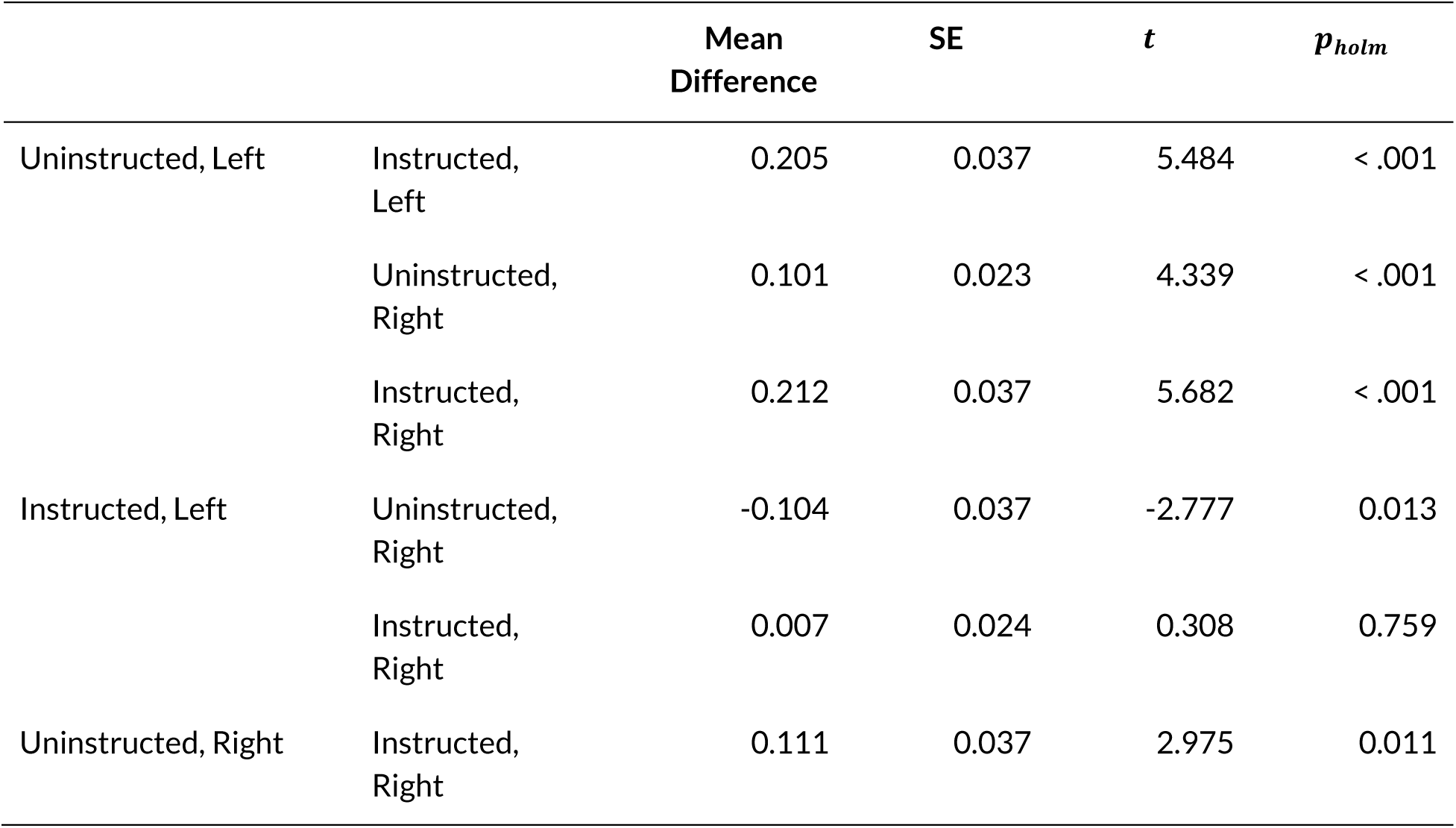
Post Hoc Comparisons - instructions ✻ Block for high lapse.

## Bibliography

Akrami, A., Kopec, C. D., Diamond, M. E., & Brody, C. D. (2018). Posterior parietal cortex represents sensory history and mediates its effects on behaviour. Nature, 554, 368–372. 10.1038/nature25510

Amjad, J., Hiratani, N., & Latham, P. (2026). Internal state is at least two dimensional. Research Square. 10.21203/rs.3.rs-7749081/v1

Andre, A., Mahato, S. P., Arora, V., Balaji, K., Lachi, D., Krishna, N. H., Xiao, J., Zhang, Y., Mao, X., Ma, W., Yu, H., Laboratory, I. B., Birman, D., Bonacchi, N., Chapuis, G. A., Catarino, J. A., Davatolhagh, F., Faulkner, M., Freitas-Silva, L., … Dyer, E. L. (2026). BrainWideBench: Benchmarking large-scale pretraining and across-animal transfer in multi-region neural recordings (arXiv:2609.22064). arXiv. 10.48550/arXiv.2609.22064

Ashwood, Z. C., Roy, N. A., Stone, I. R., Urai, A. E., Churchland, A. K., Pouget, A., & Pillow, J. W. (2022). Mice alternate between discrete strategies during perceptual decision-making. Nature Neuroscience, 25(2), Article 2. 10.1038/s41593-021-01007-z

Barron, H. C., Mars, R. B., Dupret, D., Lerch, J. P., & Sampaio-Baptista, C. (2021). Cross-species neuroscience: Closing the explanatory gap. Philosophical Transactions of the Royal Society B: Biological Sciences, 376(1815), 20190633. 10.1098/rstb.2019.0633

Bévalot, C., & Meyniel, F. (2024). A dissociation between the use of implicit and explicit priors in perceptual inference. Communications Psychology, 2(1), 111. 10.1038/s44271-024-00162-w

Bonacchi, N., Chapuis, G. A., Churchland, A. K., DeWitt, E. E. J., Faulkner, M., Harris, K. D., Huntenburg, J. M., Hunter, M., Laranjeira, I. C., Rossant, C., Sasaki, M., Schartner, M. M., Shen, S., Steinmetz, N. A., Walker, E. Y., West, S. J., Winter, O., & Wells, M. J. (2023). A modular architecture for organizing, processing and sharing neurophysiology data. Nature Methods, 20(3), Article 3. 10.1038/s41592-022-01742-6

Bruijns, S. A., Bougrova, K., Laranjeira, I. C., Lau, P. Y. P., Meijer, G. T., Miska, N. J., Noel, J.-P., Pan-Vazquez, A., Roth, N., Socha, K. Z., Urai, A. E., & Dayan, P. (2026). Infinite hidden Markov models can dissect the complexities of learning. Nature Neuroscience, 29(1), 186–194. 10.1038/s41593-025-02130-x

Brunton, B. W., Botvinick, M. M., & Brody, C. D. (2013). Rats and Humans Can Optimally Accumulate Evidence for Decision-Making. Science, 340(6128), 95–98. 10.1126/science.1233912

Burgess, C. P., Lak, A., Steinmetz, N. A., Zatka-Haas, P., Reddy, C. B., Jacobs, E. A. K., Linden, J. F., Paton, J. J., Ranson, A., Schröder, S., Soares, S., Wells, M. J., Wool, L. E., Harris, K. D., & Carandini, M. (2017). High-Yield Methods for Accurate Two-Alternative Visual Psychophysics in Head-Fixed Mice. Cell Reports, 20(10), 2513–2524. 10.1016/j.celrep.2017.08.047

Carandini, M., & Churchland, A. K. (2013). Probing perceptual decisions in rodents. Nature Neuroscience, 16(7), 824–831. 10.1038/nn.3410

Chakravarty, S., Delgado-Sallent, C., Kane, G. A., Xia, H., Do, Q. H., Senne, R. A., & Scott, B. B. (2025). A cross-species framework for investigating perceptual evidence accumulation. eLife, 14. 10.7554/eLife.105116.1

Cole, M. W., Laurent, P., & Stocco, A. (2013). Rapid instructed task learning: A new window into the human brain’s unique capacity for flexible cognitive control. *Cognitive, Affective*, & Behavioral Neuroscience, 13(1), 1–22. 10.3758/s13415-012-0125-7

Ferguson, L. A., Soukupova, M., Bouret, S., Palminteri, S., & Parkes, S. L. (2026). Reference-point dependent reinforcement learning in humans and rats. Nature Communications, 17(1), 6934. 10.1038/s41467-026-73623-x

Fernandez, K., Fengler, A., & Urai, A. E. (2025). Choice history biases evidence accumulation: A cross-species comparison from humans to mice (p. 2025.08.22.671879). bioRxiv. 10.1101/2025.08.22.671879

Findling, C., Hubert, F., Acerbi, L., Benson, B., Benson, J., Birman, D., Bonacchi, N., Buchanan, E. K., Bruijns, S., Carandini, M., Catarino, J. A., Chapuis, G. A., Churchland, A. K., Dan, Y., Davatolhagh, F., DeWitt, E. E. J., Engel, T. A., Fabbri, M., Faulkner, M. A., … Pouget, A. (2025). Brain-wide representations of prior information in mouse decision-making. Nature, 645(8079), 192–200. 10.1038/s41586-025-09226-1

Forsgren, M., Juslin, P., & van den Berg, R. (2025). Further perceptions of probability: Accurate, stepwise updating is contingent on prior information about the task and the response mode. Psychonomic Bulletin & Review, 32(3), 1284–1296. 10.3758/s13423-024-02604-2

Freeman, J. B., & Ambady, N. (2010). MouseTracker: Software for studying real-time mental processing using a computer mouse-tracking method. Behavior Research Methods, 42(1), 226–241. 10.3758/BRM.42.1.226

Garcia, B., Cerrotti, F., & Palminteri, S. (2021). The description–experience gap: A challenge for the neuroeconomics of decision-making under uncertainty. Philosophical Transactions of the Royal Society B: Biological Sciences, 376(1819), 20190665. 10.1098/rstb.2019.0665

Ghani, N. (2025). Mice wiggle a wheel to boost the salience of low visual contrast stimuli (p. 2025.08.27.672629). bioRxiv. 10.1101/2025.08.27.672629

Glasser, M. F., Smith, S. M., Marcus, D. S., Andersson, J. L. R., Auerbach, E. J., Behrens, T. E. J., Coalson, T. S., Harms, M. P., Jenkinson, M., Moeller, S., Robinson, E. C., Sotiropoulos, S. N., Xu, J., Yacoub, E., Ugurbil, K., & Van Essen, D. C. (2016). The Human Connectome Project’s neuroimaging approach. Nature Neuroscience, 19(9), 1175–1187. 10.1038/nn.4361

Glukhova, M., Tlaie, A., Taylor, R., Ferracci, P.-A., Shapcott, K., Mert, B., Arne, O., Ciuparu, A., Muresan, R. C., Havenith, M. N., & Schölvinck, M. L. (2026). Sharing the spotlight: Uncovering common attentional dynamics across species. PLOS Computational Biology, 22(4), e1014191. 10.1371/journal.pcbi.1014191

Gomez-Marin, A., & Ghazanfar, A. A. (2019). The Life of Behavior. Neuron, 104(1), 25–36. 10.1016/j.neuron.2019.09.017

Guo, Z. V., Hires, S. A., Li, N., O’Connor, D. H., Komiyama, T., Ophir, E., Huber, D., Bonardi, C., Morandell, K., Gutnisky, D., Peron, S., Xu, N., Cox, J., & Svoboda, K. (2014). Procedures for Behavioral Experiments in Head-Fixed Mice. PLoS ONE, 9(2), e88678. 10.1371/journal.pone.0088678

Hanks, T. D., & Summerfield, C. (2017). Perceptual Decision Making in Rodents, Monkeys, and Humans. Neuron, 93(1), 15–31. 10.1016/j.neuron.2016.12.003

Henrich, J., Heine, S. J., & Norenzayan, A. (2010). The weirdest people in the world? Behavioral and Brain Sciences, 33(2–3), 61–83. 10.1017/S0140525X0999152X

Hertwig, R., & Erev, I. (2009). The description–experience gap in risky choice. Trends in Cognitive Sciences, 13(12), 517–523. 10.1016/j.tics.2009.09.004

International Brain Laboratory, Aguillon, V., Angelaki, D., Bayer, H. M., Bonacchi, N., Carandini, M., Cazettes, F., Churchland, A. K., Chapuis, G., Dan, Y., Dewitt, E., Faulkner, M., Hamish, F., Haetzel, L., Hausser, M., Hofer, S., Hu, F., Khanal, A., Krasniak, C., … Zador, A. (2021). Standardized and reproducible decision-making in mice. eLife, 63711. 10.1101/2020.01.17.909838

International Brain Laboratory, Benson, B., Benson, J., Birman, D., Arlandis, J., Bonacchi, N., Bougrova, K., Bruijns, S. A., Carandini, M., Catarino, J. A., Chapuis, G. A., Churchland, A. K., Dan, Y., Davatolhagh, F., Dayan, P., DeWitt, E. E., Engel, T. A., Fabbri, M., Faulkner, M., … Witten, I. B. (2025). A brain-wide map of neural activity during complex behaviour. Nature, 645(8079), 177–191. 10.1038/s41586-025-09235-0

Johnson, P., Nieuwenhuis, S., Mejias, J. F., & Urai, A. E. (2025). A dynamical systems model of arousal-driven behavioural state transitions (p. 2025.10.31.685593). bioRxiv. 10.1101/2025.10.31.685593

Krakauer, J. W., Ghazanfar, A. A., Gomez-Marin, A., MacIver, M. A., & Poeppel, D. (2017). Neuroscience Needs Behavior: Correcting a Reductionist Bias. Neuron, 93(3), 480–490. 10.1016/j.neuron.2016.12.041

Li, J., Delgado, M. R., & Phelps, E. A. (2011). How instructed knowledge modulates the neural systems of reward learning. Proceedings of the National Academy of Sciences, 108(1), 55–60. 10.1073/pnas.1014938108

Mohammadi, Z., Ashwood, Z. C., & Pillow, J. W. (2025). Identifying the factors governing internal state switches during nonstationary sensory decision-making. Nature Communications, 16(1), 11684. 10.1038/s41467-025-66738-0

Musall, S., Urai, A. E., Sussillo, D., & Churchland, A. K. (2019). Harnessing behavioral diversity to understand neural computations for cognition. Current Opinion in Neurobiology, 58, 229–238. 10.1016/j.conb.2019.09.011

Norton, E. H., Acerbi, L., Ma, W. J., & Landy, M. S. (2019). Human online adaptation to changes in prior probability. PLOS Computational Biology, 15(7), e1006681. 10.1371/journal.pcbi.1006681

Nuiten, S. A., Lohuis, M. N. O., Schaub, A. C., Gaal, S. van, Olcese, U., Pennartz, C. M. A., Sterzer, P., & Gee, J. W. de. (2026). Baseline activity of V1 interneurons connects pupil-linked arousal to engaged behavioral state (p. 2026.02.05.704022). bioRxiv. 10.64898/2026.02.05.704022

Pisupati, S., Chartarifsky-Lynn, L., Khanal, A., & Churchland, A. K. (2021). Lapses in perceptual decisions reflect exploration. eLife, 10, e55490. 10.7554/eLife.55490

Prins, N. (2012). The psychometric function: The lapse rate revisited. Journal of Vision, 12(6). 10.1167/12.6.25

Rahnev, D., Desender, K., Lee, A. L. F., Adler, W. T., Aguilar-Lleyda, D., Akdoğan, B., Arbuzova, P., Atlas, L. Y., Balcı, F., Bang, J. W., Bègue, I., Birney, D. P., Brady, T. F., Calder-Travis, J., Chetverikov, A., Clark, T. K., Davranche, K., Denison, R. N., Dildine, T. C., … Zylberberg, A. (2020). The Confidence Database. Nature Human Behaviour, 4(3), Article 3. 10.1038/s41562-019-0813-1

Resulaj, A., Kiani, R., Wolpert, D. M., & Shadlen, M. N. (2009). Changes of mind in decision-making. Nature, 461(7261), 263–266. 10.1038/nature08275

Roy, N. A., Bak, J. H., Akrami, A., Brody, C. D., & Pillow, J. W. (2021). Extracting the dynamics of behavior in sensory decision-making experiments. Neuron, 109(4), 597–610.e6. 10.1016/j.neuron.2020.12.004

Schnell, A. E., Leemans, M., Vinken, K., & Op de Beeck, H. (2023). A computationally informed comparison between the strategies of rodents and humans in visual object recognition. eLife, 12, RP87719. 10.7554/eLife.87719

Song, J.-H., & Nakayama, K. (2009). Hidden cognitive states revealed in choice reaching tasks. Trends in Cognitive Sciences, 13(8), 360–366. 10.1016/j.tics.2009.04.009

Soric, L. (2026). Tracking Behavioral and Physiological Dynamics in Perceptual Decisions during Sustained Attention. https://studenttheses.universiteitleiden.nl/handle/1887/4303903/

Steinmetz, N. A., Zatka-Haas, P., Carandini, M., & Harris, K. D. (2019). Distributed coding of choice, action and engagement across the mouse brain. Nature, 576(7786), Article 7786. 10.1038/s41586-019-1787-x

Thura, D., & Cisek, P. (2014). Deliberation and Commitment in the Premotor and Primary Motor Cortex during Dynamic Decision Making. Neuron, 81(6), 1401–1416. 10.1016/j.neuron.2014.01.031

Thura, D., Haith, A. M., Derosiere, G., & Duque, J. (2025). The integrated control of decision and movement vigor. Trends in Cognitive Sciences, 29(12), 1146–1157. 10.1016/j.tics.2025.07.005

Urai, A. E. (2025). Structure uncovered: Understanding temporal variability in perceptual decision-making. Trends in Cognitive Sciences, 0(0). 10.1016/j.tics.2025.06.003

Urai, A. E., de Gee, J. W., Tsetsos, K., & Donner, T. H. (2019). Choice history biases subsequent evidence accumulation. eLife, 8, e46331. 10.7554/eLife.46331

Urai, A. E., Doiron, B., Leifer, A. M., & Churchland, A. K. (2022). Large-scale neural recordings call for new insights to link brain and behavior. Nature Neuroscience, 1–9. 10.1038/s41593-021-00980-9

Vloeberghs, R., Tuerlinckx, F., Urai, A. E., & Desender, K. (2026). Explaining temporally clustered errors with an autocorrelated Drift Diffusion Model (p. 2026.03.20.713186). bioRxiv. 10.64898/2026.03.20.713186

Wichmann, F. A., & Hill, N. J. (2001). The psychometric function: I. Fitting, sampling, and goodness of fit. Perception & Psychophysics, 63(8), 1293–1313. 10.3758/BF03194544

Wispinski, N. J., Gallivan, J. P., & Chapman, C. S. (2020). Models, movements, and minds: Bridging the gap between decision making and action. Annals of the New York Academy of Sciences, 1464(1), 30–51. 10.1111/nyas.13973

Yin, C., & Hiratani, N. (2026). Cortical neural landscape captures mouse-to-mouse variability in anticipatory vs. Inattentive decision making (p. 2025.07.11.664420). bioRxiv. 10.1101/2025.07.11.664420

